# PlanktonLake-CEREEP-A Freshwater Plankton Image Dataset with Semi-Automated Label Cleaning

**DOI:** 10.64898/2026.09.24.753096

**Authors:** Léo Déchaumet, Carine Puppo, David Carmignac, Samy Blusseau, Beatriz Marcotegui, Etienne Decencière, Gérard Lacroix, Jean-François Le Galliard

## Abstract

Plankton plays a fundamental role in aquatic ecosystems, influencing biogeochemical cycles and serving as a key food source for many organisms. Recent high-throughput imaging technologies enable the rapid acquisition of large volumes of microscopic images, creating new opportunities for monitoring planktonic ecosystems. However, the manual processing and annotation of the vast amounts of data generated by these devices remain time-consuming tasks. In this context, machine learning–based classification models offer a promising solution. In this data paper, we introduce a new labeled freshwater plankton dataset comprising approximately 88,000 images distributed across 43 taxa. We also present the labeling assistance method we used to facilitate dataset annotation. Finally, we present a baseline based on a convolutional neural network (CNN), which achieves a classification accuracy of 93% on our dataset.

## 1. Introduction

Plankton encompasses a diverse group of aquatic organisms that drift with currents and lack the ability to swim against them. This broad definition encompasses an exceptional diversity and size distribution of organisms, ranging from microscopic microalgae to giant deep-sea siphonophores, as well as larval stages and small crustaceans such as copepods, cladocers or krill, a major food source for aquatic predators such as fishes, birds and mammals. Planktonic organisms play fundamental roles in ecosystems, and are involved in multiple global biogeochemical cycles, including the carbon cycle (Worden et al., 2015). As the foundation of aquatic food webs at the surface of oceans and in freshwater ecosystems, phytoplanktonic organisms produce organic matter by photosynthesis (Field et al., 1998) which in turn feeds herbivorous zooplankton, such as small crustaceans. These organisms are then eaten by larger animals, including predatory zooplankton, allowing energy to move up the food chain and supporting major ecological services such as provisioning of food. Even if freshwater ecosystems occupy only 0.8 % of the total Earth land surface, they support almost 6% of all the described species worldwide and have large significance for terrestrial ecosystems and human societies (Declerck and Senerpont Domis, 2022; Dudgeon et al., 2006). Research on freshwater plankton ecosystems has been extremely fruitful to document its taxonomic diversity, its numerous functional roles and its sensitivity to global changes in lakes and rivers worldwide (Woodward et al., 2010). For instance, zooplankton can serve as a bioindicator of freshwater ecosystem health and water quality. Rotifers, for example, can be used to monitor the impact of toxicants such as endocrine disruptors or nanoparticles (Snell and Marcial, 2017). Quantitative information on plankton abundance, diversity and size distribution is also useful to model trophic webs or networks and therefore predict the future of freshwater ecosystems (Declerck and Senerpont Domis, 2022).

The study of freshwater plankton, particularly zooplankton (the heterotrophic fraction of plankton), poses numerous challenges due to the taxonomic and functional diversity, morphological heterogeneity and very wide size range of this group of organisms. Classical sampling techniques use size sorting protocols together with manual observation by microscopy and classification by experts (Stanislawczyk et al., 2017), or more recently molecular biology techniques such as metabarcoding. However, these sampling methods present drawbacks. DNA-based methods require large and high quality reference databases, and may not be well suited for quantitative analysis without a proper calibration that can depend on several details of the sampling and bioinformatic protocols (Xiong et al., 2020). Besides that, manual observation of zooplankton using a microscope is very accurate but can be very time consuming, might be prone to errors without proper expert guidance and might not allow to sample efficiently rare species, as only a limited number of images can typically be examined. Thus, new automated imaging tools such as the FlowCam (Sieracki et al., 1998) or the Zooscan (Gorsky et al., 2010) have been developed to overcome these problems and facilitate the process by allowing the acquisition of hundreds of microscopic images per second. However, imaging methods still require human annotation, and assigning a taxonomic label to each image remains a time-consuming and error-prone task.

Most of the software associated with imaging tools extract morphological features of the objects present in the images, such as area, biovolume, and circularity. These features can be used to facilitate the work of the annotator. For example, the “Visual Spreadsheet”, a proprietary tool associated with the FlowCam, allows the user to define rule-based filters for each species based on these morphological traits. The web platform Ecotaxa (Irisson et al., 2022), widely adopted by the plankton research community, provides a machine learning tool based on random forests to assist annotation. However, these methods are often less accurate than modern deep learning approaches for plankton classification, especially for rare planktonic classes (Panaïotis et al., 2026). Indeed, deep learning based methods such as Convolutional Neural Networks (CNNs) or vision transformers have become dominant approaches for image classification tasks, due to their ability to learn relevant visual features directly from raw images. In the context of plankton imaging, CNN-based models have demonstrated strong performance, particularly on large-scale marine datasets where annotated data is relatively abundant (Panaïotis et al., 2026). However, such success is strongly dependent on the availability of large and high-quality labeled datasets. Several reference datasets exist for marine plankton, but freshwater plankton datasets remain significantly underrepresented. This lack of datasets limits the development of robust image classification models in freshwater ecosystems.

In this work, we address this gap by constructing PlanktonLake-CEREEP, a freshwater plankton image dataset labeled by experts, with a particular attention given to label quality for future reuse. We first review the state-of-the art freshwater plankton datasets and automated classification methods tested in previous studies. Then we detail the computer-assisted, semi-automatic label-cleaning strategy we applied to detect and correct efficiently annotation errors, a strategy that is designed to be easily reproducible across different datasets. To facilitate its adoption by the community, and make the workflow accessible for non-expert users, we provide an easy-to-use Python package and a graphical interface. We apply this framework to a large image dataset from artificial lakes and compare the efficiency of our classification model with a previously published model by Walter et al., 2025.

## 2. Summary of previous works

### 2.1. Freshwhater Plankton Datasets

We reviewed the literature and a global database of plankton images for published datasets of freshwater plankton images. Our literature review was performed in a non-systematic manner with Google Scholar and we explored datasets in Ecotaxa repository in March 2026 (https://ecotaxa.obs-vlfr.fr/). Ecotaxa is a widely used platform for publishing plankton image datasets (Irisson et al., 2022), and this repository hosts over 700 million images from approximately 900 institutions. However, most of the datasets focus on marine plankton images. As far as we know, there is no simple way to filter for freshwater plankton dataset via the online interface, but we performed a search on the datasets names using the keywords “lake” (as well as “lac” and “lago”), “freshwater”, “pond”, and “river”, which yielded 12 projects containing more than 1000 validated images. In addition to Ecotaxa, we found three datasets published independently from Ecotaxa and associated with their corresponding data papers. This includes the ZooLake dataset (Kyathanahally et al., 2021b) which consists of 17900 plankton images from the Lake Greifensee in Switzerland. These images were acquired using the Dual Scripps Plankton Camera and annotated manually by experts. As a baseline, the authors built a machine learning classification model combining handcrafted features extracted from the images and deep learning features. The FREPJ (Otake et al., 2024) is a dataset composed of plankton images taken from 87 Japanese lakes and reservoirs to provide a reference for the development of automated classification models. Images were acquired using a microscope at magnification of *×*40 and *×*100 and were also annotated manually by multiple experts. The last dataset is PlanktonFlow (Walter et al., 2025), which contains images taken with the FlowCam 8100 and FlowCam Macro in samples from aquatic mesocosms located in Rennes, France. In this dataset, images were also annotated manually by multiple experts with the possibility of annotation errors specifically acknowledged. A summary of the existing datasets is provided in table 1.

**Table 1.**
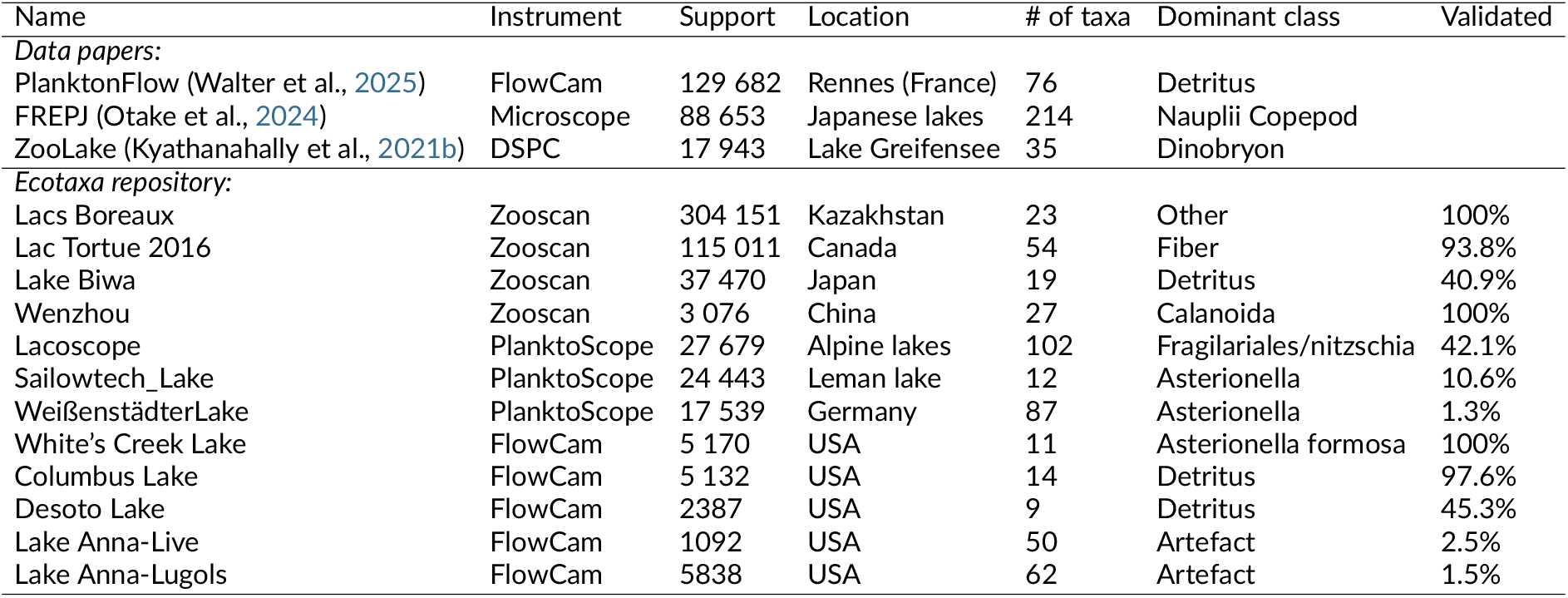
Public freshwater plankton datasets. All datasets, except the first three, are stored in Ecotaxa (https://ecotaxa.obs-vlfr.fr/). Support denotes the number of annotated images.

### 2.2 Image Classification

Traditional image classification models, such as Support Vector Machines (Cortes and Vapnik, 1995) or Random Forests (Breiman, 2001), rely on handcrafted features, such as particle area, measures of circularity, mean intensity, and more complex features like Hu moments (Hu, 1962) or Fourier descriptors (Zahn and Roskies, 1972). Some software can also infer the biovolume of plankton from images, which is an important feature in analyses of trophic networks (Gorsky et al., 2010; Sieracki et al., 1998). For example, a tool like the Visual Spreadsheet, associated with the FlowCam, subtracts the background of the image to segment the object and extract shape features. However, the extracted features are usually not very robust, as they can be affected by image quality, the presence of multiple objects or detritus, and even the software version used to extract these characteristics. Moreover, these features may struggle to discriminate efficiently between some closely related and morphologically similar taxa, as they are not always able to capture the complex morphology of plankton organisms. For these reasons, traditional machine learning models based on handcrafted features may have limited performance in plankton image classification tasks (Panaïotis et al., 2026).

Alternatively, deep learning–based methods such as Convolutional Neural Networks (CNNs) are powerful approaches for image classification tasks, due to their ability to learn relevant visual features directly from raw images instead than from pre-defined features (LeCun et al., 2015). This technique has been widely used in marine plankton classification, where it can achieve very high classification performance (Dai et al., 2016; Guo and Guan, 2021; Hassan et al., 2025; Kerr et al., 2020; Lee et al., 2016; Lumini and Nanni, 2019; Orenstein and Beijbom, 2017). The scientific literature focusing specifically on freshwater plankton images includes three studies. The previously cited ZooLake data paper (Kyathanahally et al., 2021b) proposed a baseline classification model that combines handcrafted features and deep learning methods with high classification success and the ability to outperform previously used models on other freely available datasets. Ito et al. (Ito et al., 2023) used a hierarchical attention layer to learn the hierarchical structure inherent to taxonomic labels. Recently, Walter et al. (Walter et al., 2025) proposed an easy-to-use deep learning pipeline to classify lake plankton acquired with FlowCam and reached high classification success with optimized architectures, yet limited performances in taxa with a high intra-class variability.

One important limitation of deep learning methods is that they are computationally more demanding than classical methods such as random forests. The massive volume of data collected by modern imaging tools with high acquisition rates, such as the FlowCam, thus introduces challenges in terms of efficiency (Eerola et al., 2024). For example, Zimmerman et al., 2020 needed a plankton classification model small and fast enough to run in a Raspberry Pi. As CNN architectures did not fulfill these constraints, these authors chose a decision tree model in order to have the best trade-off between accuracy and inference time. It is therefore important to design models that are fast enough to process large amounts of data. The authors of TANet, a light-weight deep learning method (Li et al., 2019), show that a model can be smaller, faster in terms of inference time and still be competitive in classification performance. Finally, previous studies have shown that deep learning is becoming a major contributor of greenhouse gases emissions, thus it is important to monitor energy consumption and carbon footprint of machine learning models. However, these metrics are rarely reported in the machine learning literature. Henderson et al., 2020 analyzed 100 papers randomly taken from the prestigious NeurIPS conference, and noticed that only one paper reported the energy consumption of its model, and 0 estimated its carbon footprint.

### 2.3. Labeling Quality

The performance of a machine learning classification model relies strongly on the quality of the original labels. However, in many cases, both resources and the number of experts available for image annotation are limited, and even experts may make mistakes. For example, Culverhouse et al., 2003 analyzed the performance of dinoflagellates identification approaches and showed that even with multiple experts, a significant number of mislabeled samples can remain in the dataset after the annotation process. These labeling errors can impact a classification model: if the test set contains a significant proportion of labeling errors, the reliability of the evaluation is compromised. Labeling errors in the training set can also degrade the learning process, as deep learning models often possess sufficient expressiveness to overfit noisy labels present in the data (Zhang et al., 2021).

Several studies have addressed the challenge of automatically detecting potentially mislabeled samples in image datasets in order to reduce mistakes in the training or test sets. These methods typically involve training a machine learning model on a potentially noisy dataset and identifying suspicious samples by detecting inconsistencies between the model’s predictions and the provided labels. For instance, Bahri et al., 2020 applied a k-nearest neighbors (KNN) algorithm on the features produced by a trained model, with the intuition that if an image has a different label than its neighbors in the feature space, it may be mislabeled. Northcutt et al., 2021 observed that the average predicted probability can vary across classes. They therefore computed class-specific thresholds to identify potentially mislabeled images.

Once suspicious samples are spotted, several options are available. A straightforward approach is to simply discard the suspicious images from the training set, thereby a large part of label noise can be removed. However, it may also eliminate inherently difficult or ambiguous samples, and oversimplify the training process. An alternative approach is to manually review the suspected samples and correct their labels when necessary. While this strategy preserves the diversity and complexity of the dataset, it requires additional human annotation effort.

## 3. Data Annotation and Semi-Automatic Label Cleaning

We present a data annotation and semi-automatic label cleaning procedure combining human and machine-made annotation methods to help produce a golden standard, reference dataset for future, automated freshwater plankton image analysis. The constitution of PlanktonLake-CEREEP involved several major steps adapted to the specificity of our image acquisition method and to the number and quality of images, but our method is sufficiently general to suit similar use cases.

In particular, it is adapted to situations where taxonomic expertise is limited since it uses machine-made annotations to improve incorrect human-made classifications in an iterative manner.

For clarity, each version of the dataset was assigned a different name. The first version V1.0 contained 58103 images and 26 taxa. In December 2025, new labeled images were added to our dataset, to raise the number of images from rare taxa, add new ones and create V2.0 of the dataset. From this version, a test set was created, the procedure is explained in Section 3.4. For the train set only, a second cleaning phase was performed (Section 3.5) to produce the final version, called V2.1. The dataset is available on Zenodo (https://zenodo.org/records/22012197), while the source code is available on GitHub (https://github.com/ixalodecte/PlanktonLake). The general procedure is explained in Figure 1 from image acquisition to dataset production.

**Figure 1.**
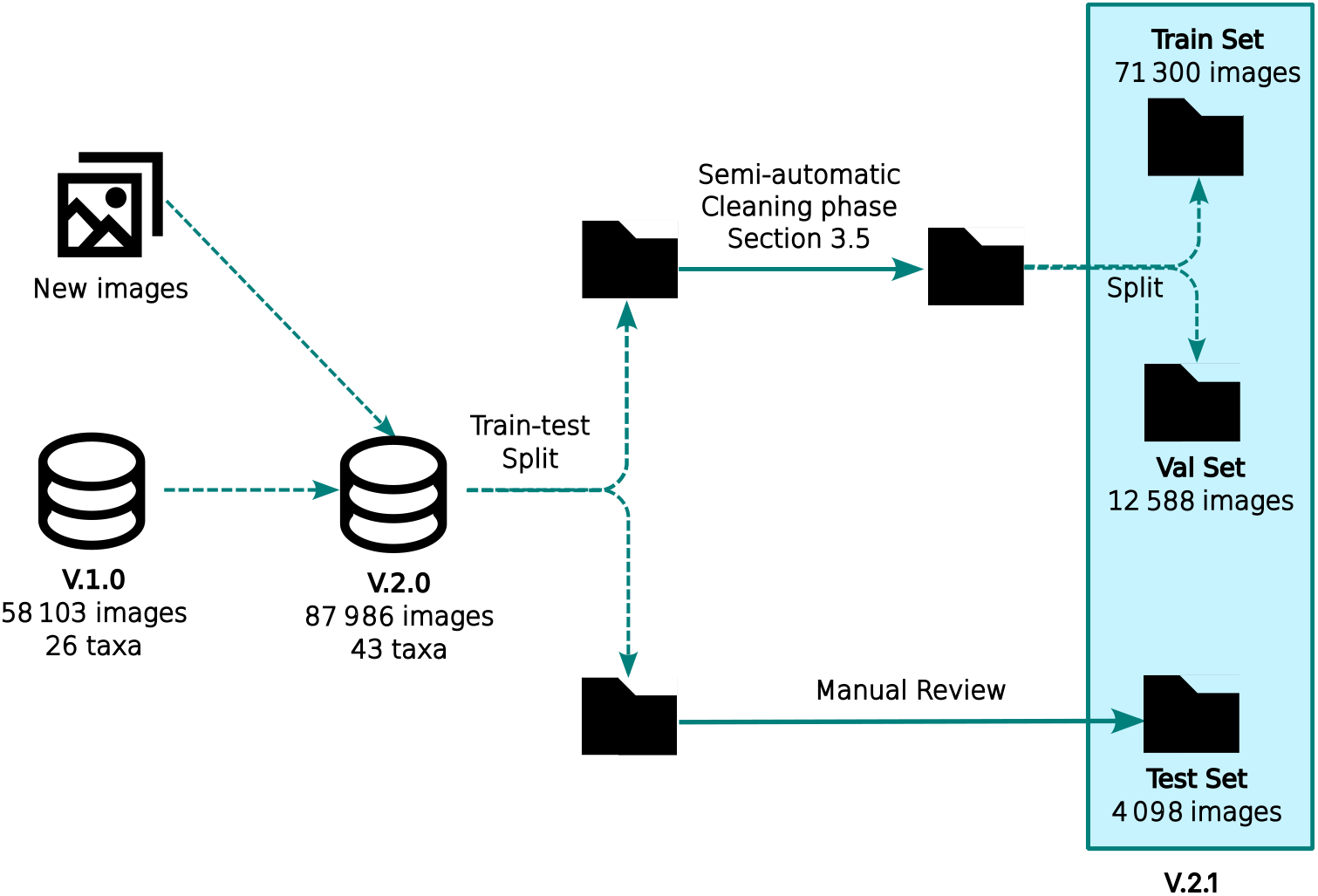
The initial dataset was labeled following the procedure described in section This dataset was reviewed using a model-based approach that identified samples with discrepancies between the assigned labels and model predictions, followed by expert validation and correction of annotation errors, resulting in dataset version V1.0. In December 2025, new labeled images were added to our dataset, to raise the number of images from rare taxa, add new ones and create V2.0 of the dataset. From this version, a test set was created, the procedure is explained in Section 3.4. For the train set only, a semi-automatic cleaning phase was performed (Section 3.5) to produce the final version, called V2.1.

**Figure 2.**
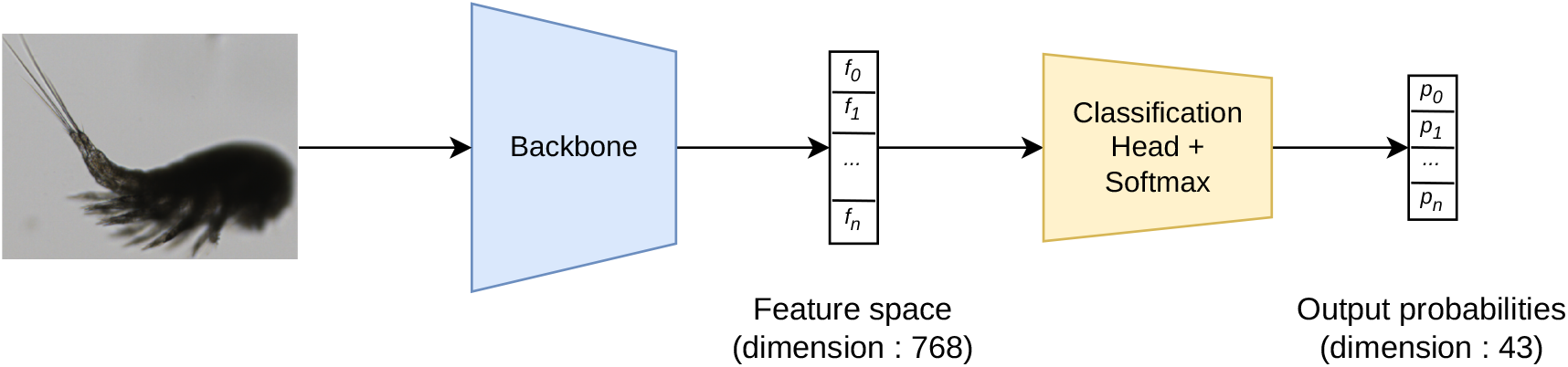
Architecture of a classification model. The backbone is a convolutional neural network that learns a projection of the input image into a feature representation space of a given dimension (768 for ConvNeXt). The classification module, consisting of a fully connected layer, takes as input the feature vector encoding the image and outputs the probability of the image belonging to each class.

### 3.1 Image acquisition

Water samples were collected on 16 experimental ponds located at the PLANAQUA facility in France about monthly between January 2015 and December 2025. These artificial ponds measured each 30 m × 15 m × 3 m with a water volume of 750 m3, oriented North to South and arranged in a grid (Schmidt et al., 2024). Each pond contains a 3 m deep central pelagic zone and outer shallow littoral zones (0.25 m deep) containing macrophytes. Plankton was collected in the central, pelagic zone during long-term experiments (Schmidt et al., 2024). Zooplankton and large phytoplankton specimens were concentrated in the field by filtering 90 liters of water (fifteen spatial points and three depths at each point) through a 50*µm* mesh netting. During the sample filtering, organisms are initially retained on the filter. Over time, as larger organisms accumulate on its surface, they progressively clog the filter, preventing further passage of particles through it. Then, the samples were fixed and preserved with a 99% ethanol solution until processing in the laboratory. FlowCAM 8000 imaging cytometer equipped with VisualSpreadsheet software version 6.0.2.153 (Yokogawa Fluid Imaging Technologies, Inc.) was used to analyse plankton samples. Samples collected with the 50*µm* mesh net were split into two subsamples using two mesh nets (850*µm* and 400*µm*) in order to separate small and large particles. The 850-400*µm* subsample and *<* 400*µm* subsample (*≈* 100mL of 96% ethanol) were diluted to 500mL with deionized water (into a 500mL bottle), and homogenized using a stirring barrel. Before each sample analysis, the prime system settings were used to flush the flow assembles. This was performed six times before each sample analysis followed by filling the assembly with deionized water to remove all particles from the previous sample. Then, the manual prime was used to draw each sample into the flow cell. For the 850*µm*-400*µm* subsample, the FlowCAM was set to a flow rate of 4 mL/min and 3000 particles were recorded in autoimage mode using combinations of 2x objective with the 1000*µm* flow cell and the 12.5mL syringe. For the *<*400*µm* subsample, we recorded in autoimage mode using combinations of 4x objective with the 600µm flow cell and the 5mL syringe. The FlowCAM was set to a flow rate of 3 mL/min and three technical subsamples were made. We used the ESD (Equivalent Spherical Diameter) as filter. The first technical subsampling included particles between 200-3000*µm* with 1000 particles were recorded. The second technical subsampling included particles between 100-200*µm* and 1000 particles were recorded . The last technical subsampling included particles between 20-100*µm* and 2000 particles were recorded. This protocol was necessary to sample efficiently across the size range of particles, which is dominated by small organisms and detritus.

### 3.2 Annotation Procedure

For identification, classification, and quantification of all the plankton particles, the Visual-Spreadsheet software version 6 (Version: 6.0.2.153) Yokogawa Fluid Imaging Technologies, Inc.) was used. To facilitate the annotation process, the user can define filters (particle area, biovolume, etc), to sort the data. However, this process lacks precision and contains a significant number of errors, so the “pre-sorting” is not very accurate and requires manual validation of every image.

#### Duplicates

It may happen that the Visual Spreadsheet software saves the same image multiple times. These copies appear in the dataset as *strict duplicates*, and they are strictly identical. To detect and remove these duplicates, the SHA-256 hash of each image is computed. For images that share the same hash, only one instance is retained.

#### Taxonomy

EcoTaxa recently transitioned to the World Register of Marine Species (WoRMS) taxonomy (marinespecies.org) (WoRMS Editorial Board, 2026). Given that WoRMS constitutes a reference framework for plankton classification, this taxonomy is adopted for our dataset. The labels given by the experts can be found at different levels of the taxonomic hierarchy. For example, “Cyclopoida” is an order, “*Daphnia*” is a Genus and “Dinoflagellata” is an Infraphylum. The level of precision depends on both the annotator’s expertise and the taxa. For instance, species within the genus *Keratella* (e.g., *Keratella cochlearis, Keratella quadrata*) are relatively easy to recognize, which is not the case for species of Cyclopoida. Alongside plankton images, our dataset also contains detritus, and unidentifiable blurry images. These images represent approximately 70% of our dataset, they are grouped into a dedicated class labeled “Unknown.”

We generated a csv file that contains, for each label, different taxonomy levels: Superdomain (which can take only 2 values, Biota or Unknown), Kingdom, Phylum, Class, Order, Family, Genus and Species. In addition, intermediate levels of classification are provided (for example, Super-class or Infraorder), but these intermediate levels can be missing for some labels (marked as “-” in the csv file). The script used to retrieve the full taxonomy from a taxon using the WoRMS API is provided with the code.

#### Multiple objects in the image

While the vast majority of images contain a single element, some images may include multiple objects. This is often the case when those objects are very close to each other, and the software fails to segment individual parts. Several situations may occur. First, multiple individuals of the **same taxon** may be present in the image. This frequently occurs with *Ceratium* and *Keratella cochlearis*, whose morphology tends to promote grouping. In such cases, the image is labeled in the same way as one containing a single individual, using the corresponding taxon label. Second, a **detritus** may be present alongside an organism. In this situation, the image is annotated with the corresponding plankton taxon. Finally, **multiple plankton taxa** may appear in the same image. If this situation were common in the dataset, it would require a multilabel annotation approach. However, as shown in section 3.4, such cases are very rare. Therefore, a “one image–one taxon” rule is adopted, and each image is assigned the label corresponding to the rarest species present (i.e., the least represented species in the dataset). During the second cleaning phase, however, all images detected as containing multiple species were removed from the dataset to ensure that each image unambiguously represents a single taxon (see section 3.5).

### 3.3 Automatic Classification Method

This section presents the CNN we use to detect labeling errors in section 3.5. It also serves as baseline classification model for PlanktonLake-CEREEP (section 4). The performance metrics for its evaluation are introduced at the end of this section.

#### 3.3.1. Method

For all our experiments, the ConvNeXt-tiny architecture (Liu et al., 2022) is chosen as the backbone of our model. The optimizer used is AdamW (Loshchilov and Hutter, 2017). To prevent overfitting, we use data augmentation, *i*.*e*. random transformations are applied to each input image during training. The following transformations are applied randomly, with varying intensities: Random shift of Hue, Saturation and Value, Gaussian Blur, resize, horizontal flip, vertical flip, 90-degree random rotation, random shift, scale, and rotation, zero padding, and Coarse Dropout.

Three important aspects of this transformation process need to be highlighted. First, to enable batch processing in CNNs, all images must have the same size. This can be achieved by either directly resizing the image, which alters the aspect ratio of the object (Kyathanahally et al., 2021b; Sánchez et al., 2019), or resizing the image while preserving the aspect ratio (so the longest edge matches the target size) and padding the shorter edge to create a square (Dai et al., 2016; González et al., 2019). In preliminary tests, both strategies gave equal performances. The second option was chosen arbitrarily. As discussed in the results section, different input image sizes were tested. A size of 128 *×* 128 was chosen as a good trade-off between computational performance (small image size) and classification performance. Second, even if the data acquisition process retains limited color information, data are still collected and stored as RGB images. Performances of the model using RGB vs. grayscale images were compared to assess the impact of color. The accuracy on the validation set was slightly better with RGB images, therefore this input format was used to train the model. Finally, Gaussian blur was randomly applied with a relatively high maximum intensity to align the model’s robustness to blur with the blurry images originally present in the dataset.

#### 3.3.2. Performance metrics

To evaluate the performance of our models, multiple standard metrics are used: accuracy, recall, precision and F1-score. Our models are also evaluated in terms of inference time.

##### Accuracy

Accuracy measures the proportion of correctly classified images. While being easy to understand, it is not suitable for imbalanced datasets, as the minority classes have a minor impact on this metric. In our case, the test set is more balanced than the full dataset, thus measuring accuracy makes sense.

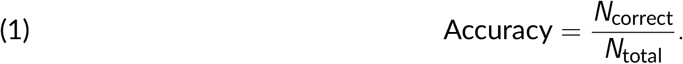

##### Precision, Recall, F1

Precision and recall are metrics used in a binary classification setting. Thus, this metric should be computed class by class. For a given class *c*, Precision_*c*_ is the proportion of samples predicted as belonging to class *c* that are actually correct. Recall_*c*_ is the proportion of samples of class *c* that are correctly identified by the model:

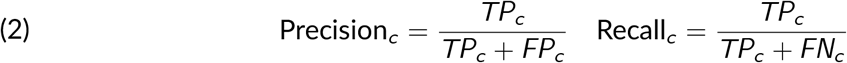

where *TP*_*c*_ is the number of true positive for class *c*, and *FN*_*c*_ and *FP*_*c*_ are the numbers of False negatives (images belonging to class c that are predicted as another class) and false positives (images from other classes that are incorrectly predicted as class *c*), respectively.

The F1 score is a metric that aggregates recall and precision as their harmonic mean:

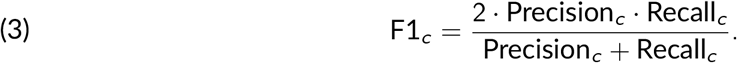

It takes values in [0, 1] and the higher it is, the better.

##### Macro metrics

Macro metrics for precision, recall and F1 are computed as the mean of each class metric. Macro metrics are useful to evaluate the performance of a model on an imbalanced dataset, because it gives equal importance to each class, including the rare ones. Macro metrics are sensitive to rare classes, and penalize the score if the model fails to classify correctly a rare class.

##### Computational efficiency and environmental impact

Alongside classification performance metrics, the computational efficiency of our model is assessed. Inference latency in milliseconds is measured and the power consumption and carbon footprint of a single training session is estimated using the CodeCarbon Python package. Inference latency is measured on a laptop equipped with an Intel CoreTM Ultra 5 225U × 14, and the carbon footprint is measured on a GPU V100.

### 3.4. Training, validation and test sets

Machine learning pipelines commonly divide a dataset into training, validation, and test sets. The training and validation sets are used to optimize the model, whereas the test set is used to evaluate its performance on previously unseen images after the training phase. First experiments on dataset V1.0 randomly divided the dataset into training and test sets. When training our classification model, a significant proportion of the model’s errors on the test set actually came from labeling errors. These errors arise from several sources. First, **taxonomic ambiguity** can make annotation difficult. While some taxa, such as *Ceratium* or *Keratella cochlearis*, are easily recognizable, others, including *Hexarthra* and *Polyarthra*, are much more difficult to distinguish. Second, in some very rare cases **multiple organisms may be present in the same image**. As discussed in the previous section, a model may correctly predict a taxon that is present in the image but not the one assigned by the annotator, thereby artificially degrading the reported performance. Finally, **image quality** can also be a limiting factor, as some images are too blurred to allow reliable organism identification (see fig. 3).

**Figure 3.**
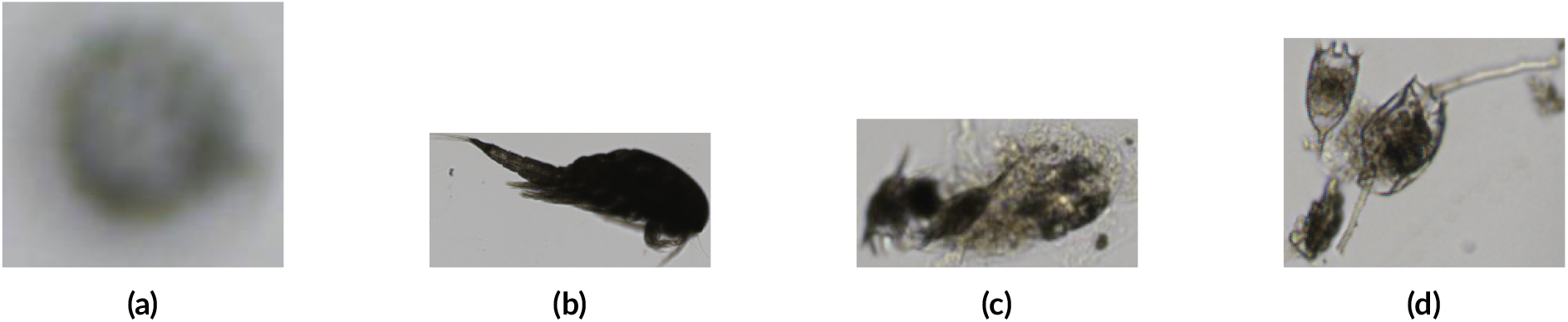
Examples of images difficult to label. a) Image with excessive blur leading to an ambiguity. It may be a *Volvox* or a *detritus*. This image was labeled as a detritus. b) A *Cope-pod* whose orientation does not allow discrimination between *Calanoid* and *Cyclopoid* groups. This image (present in the test set), is excluded). c) Two *Keratella cochlearis* individuals adjacent to detritus. (labeled as *Keratella cochlearis*) d) A *Keratella cochlearis* and a *Pompholyx*. This type of images is excluded from the test set.

**Figure 4.**
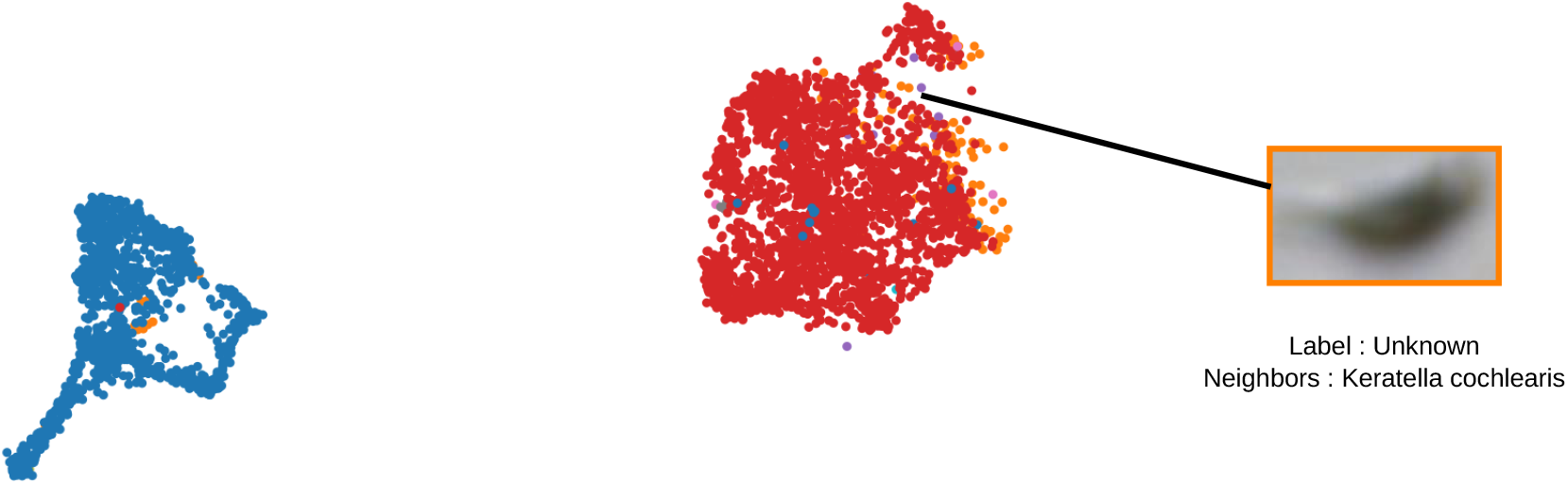
Visualization of the representation space. The UMAP algorithm is used to reduce the embedding dimensionality from 768 to 2. Each class is represented by a specific color (for example, blue points correspond to *Ceratium*, while red points correspond to *Keratella cochlearis*). Images whose nearest neighbors do not match their ground-truth label are considered suspicious. For readability, this figure shows only a cropped region of the complete UMAP embedding : not all classes are displayed.

While research has explored automatic methods for detecting suspicious annotations in datasets (see next section), the choice has been made not to apply these methods to the test set. Automated cleaning could introduce bias, as the test set would then be partially aligned with the predictive model. If a model is able to reproduce label errors in the train set, this could artificially simplify the evaluation task and lead to overly optimistic performance estimates: for all these reasons, we decided to create a “gold standard” test set, which was manually reviewed by experts. This test set had to be as small as possible to be reviewed manually, but large enough to provide a good estimation of the performance of a classification model. The test set was created following this procedure. First, for each class, 22% of the images were randomly selected, with a maximum of 250 images per class. An exception was made for the *Unknown* class: as it is highly overrepresented and heterogeneous, the test set contains 500 samples for this class. Sampling was performed to ensure a balanced distribution across the two magnification levels of the FlowCam. Second, each selected sample was reviewed individually, and its label is corrected when necessary. Dedicated folders were created to store images containing multiple plankton organisms from different species (e.g., *cyclopoide+ceratium*). Finally, in some cases, distinguishing between *Calanoids* and *Cyclopoids* is particularly challenging. Eight copepods that could not be reliably identified were found. Additionally, we found one cladoceran that could not be distinguished between *Ceriodaphnia* and *Daphnia*. These nine images were excluded from the test set. After this procedure, our test set contains 4,098 images. Among these images, only five samples containing multiple species were found. As the test set was randomly sampled from the dataset, we can reasonably estimate that the proportion of images containing multiple species in the full dataset is very low (approximately 0.1–0.2%). These five images were excluded from the test set. Overall, 163 images (3.9% of the test set) had their labels corrected during the annotation review process.

### 3.5. Semi-automatic data cleaning

While reviewing all test images is feasible, the training set is substantially larger. Therefore, an automated computer-assisted cleaning procedure is applied to detect potentially suspicious samples, which are selected for manual review. The automatic detection of labeling errors relies on the assumption that, when trained on a large enough dataset containing a majority of correctly labeled samples, a machine learning model is already quite reliable. Hence, its output when given a new sample can be taken into account to label or re-label that sample.

In order to apply this principle and automatically check the labels of a whole dataset, a 5-fold cross-validation strategy was employed. The dataset was divided into five equally sized subsets (folds). At each iteration, four folds were used to train a classification model with a fixed number of 20 epochs, while the remaining fold was never seen during training and is used for prediction. This validation fold was the one checked for possible labeling errors. This process was repeated five times to ensure that every image was evaluated exclusively by a model that had not been trained on that image.

To improve the robustness of the detection, two complementary methods were used to find suspicious samples. For both methods a label quality score between 0 and 1 was calculated for each sample: the smaller the score, the more likely the corresponding label is to be wrong. The first quality score is based on the **self confidence**. Confidence refers to the probability a model assigns to a class, for a given image. It ranges between 0 and 1, and is computed as the output of the final layer of the model (logits), processed with the softmax function. The **self confidence** method consists of looking at the confidence that a model, trained on (noisy) labels, gives for the ground truth label. A threshold of 0.3 was chosen to mark a sample as suspicious, which yielded 1532 suspicious samples. The choice of the threshold depends on the time we want to allocate to the reviewing process, as a higher threshold would result in a higher number of images to review. The second technique is based on the *k*-nearest neighbors algorithm (**KNN**) (Bahri et al., 2020) applied to the learned *representation space*. During the training process of a CNN, the model learns a representation, where each image is mapped to a feature vector. This space is expected to capture features that are relevant for the classification task. The intuition is that neighbouring samples in this space are likely to share the same label. If this is not the case, then this image may be mislabeled. We choose *k* = 11, ensuring that *k* remains smaller than the minimum class support (which is 20). This prevents minority-class samples from being systematically penalized due to an insufficient number of same-class neighbours in their local neighbourhood. Here the label quality score of a given image is defined as the proportion of nearest neighbors that have the same label as the considered image. A threshold of 0.3 is chosen for this method as well, to find 1772 suspicious samples.

The final set of suspicious samples is built as the union of the samples found by the two methods. This set contains 2208 samples 1096 of which were flagged as suspicious by both methods (more than one third, see table 2). All the suspicious samples were reviewed by an expert using a graphical user interface and were either reassigned to an existing class, retained in their original class, or identified as containing multiple species and subsequently discarded from the dataset. To sort the images in priority order, the minimum of the two scores is chosen, and the most suspicious images (those with lower scores) are reviewed first. The label distribution is shown in fig. 5. Results of this cleaning phase are shown in next section 4

**Table 2.** Performances of the two methods used for the second cleaning phase. Precision is the proportion of suspicious images that are actually mislabeled. Recall cannot be computed because the ground-truth label errors are unknown. The *union* corresponds to all images flagged as suspicious by at least one of the two methods (or by both). The *intersection* contains only the samples identified as suspicious by both methods. Finally, *knn only* and *self-confidence only* refer to the suspicious samples detected exclusively by the KNN method or the Self-Confidence method, respectively..

| Method | Number of suspects | Precision |
| --- | --- | --- |
| knn | 1772 | 41.8% |
| self-conf | 1532 | 50.5% |
| union | 2208 | 43.3% |
| intersection | 1096 | 50.9% |
| knn only | 676 | 26.9% |
| self-conf only | 436 | 49.2% |

**Figure 5.**
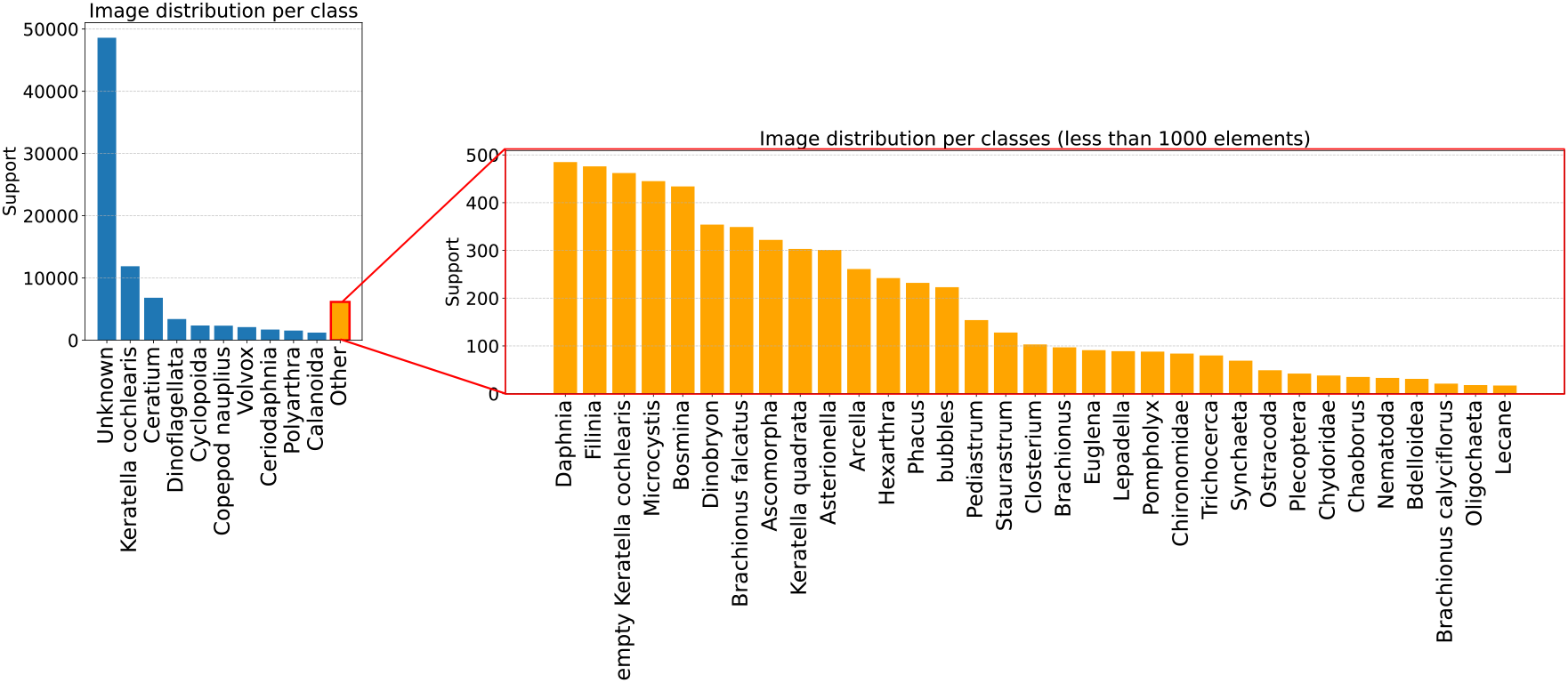
Image distribution of the final version of the dataset, V2.1.

## 4. Results

### 4.1. Efficiency of the label-cleaning

After the reviewing process, the proportion of wrong label among suspected images for both methods can be measured (table 2). This table shows that the Self-Confidence method is more precise than the KNN method on our dataset. The two methods appear to be complementary, as each detects some label errors that are not identified by the other. Therefore, combining both methods is beneficial, as it increases the overall number of detected label errors (recall).

While table 2 quantifies the precision of the automatic error detection methods, it does not provide information about the types of annotation errors that remain undetected. To better characterize these limitations, we can compare the proportions of corrected samples for each class between the fully manually reviewed test set and the semi-automatically corrected training set. For several classes, the correction rates are similar in both datasets. This is the case for *Unknown* (0.8% in the test set vs. 0.7% in the training set; 330 corrected samples out of 48,166), *Ascomorpha* (7.1% vs. 5.8%;), *Brachionus* (9.5% vs. 7.8%; ), *Chaoborus* (14.3% vs. 16.7%;), and *Closterium* (4.2% vs. 3.8%; ). The dominant label transitions are also consistent across the two datasets, including *Ascomorpha → Unknown, Unknown → empty Keratella cochlearis, Cyclopoida → Unknown, Cyclopoida → Daphnia*, and *Calanoida → Cyclopoida*.

In contrast, several classes exhibit substantially lower correction rates in the training set than in the manually reviewed test set. The largest discrepancy is observed for *Dinoflagellata*, for which only 17 out of 3,148 training samples (0.5%) were corrected, compared with 25.6% in the test set. This difference is almost entirely explained by the *Dinoflagellata → Unknown* transition, which is frequently identified during exhaustive manual review but rarely detected by the semi-automatic procedure. Similar behavior is observed for *Keratella cochlearis*, where only 147 of 11,537 training samples (1.3%) were corrected despite a correction rate of 5.6% in the test set, mainly due to missed *Keratella cochlearis → Unknown* transitions. Likewise, *Ceratium* shows a correction rate of only 0.7% in the training set (43/6,530) compared with 4.0% in the test set, again largely explained by the *Ceratium → Unknown* transition. More moderate discrepancies are also observed for *Cyclopoida, Calanoida*, and *Asterionella*. These classes are represented by hundreds to thousands of training samples, indicating that the observed differences cannot be explained by insufficient support. Instead, the results suggest that the semi-automatic procedure effectively identifies systematic annotation inconsistencies but is less sensitive to ambiguous samples that are ultimately relabeled as *Unknown*.

### 4.2. Baseline classification results

In this subsection, we present the classification results obtained on the test set of PlanktonLake-CEREEP by models trained using the cleaned version of the dataset. Results for the architecture “ConvNext-tiny” and multiple image sizes are shown in table 3. A single training of the ConvNext model with input size 128 *×* 128 emitted 0.0079 kg CO_2_. This estimate was obtained using the CodeCarbon package, which measures the energy consumption during training and estimates the associated CO_2_ emissions based on the carbon intensity of the electricity grid in France.

**Table 3.** Performance (min–max) on 5 runs, for four models trained with different input image sizes, evaluated on the test set. Models are trained with the dataset before the second label cleaning phase (V2.0) and after (V2.1)

| Dataset | Model | Accuracy | Precision | Recall | F1 | Latency |
| --- | --- | --- | --- | --- | --- | --- |
| V2.0 | ConvNext (64 × 64) | 92.8–93.4 | 92.3–93.4 | 90.6–92.3 | 91.0–92.4 | 18 ms |
|  | ConvNext (128 × 128) | 92.9–93.6 | 92.7–93.9 | 90.3–92.2 | 90.9–92.8 | 40 ms |
|  | ConvNext (224 × 224) | 92.9–93.5 | 93.5–95.3 | 89.7–91.3 | 91.0–92.7 | 86 ms |
| V2.1 | ConvNext (64 × 64) | 93.0–93.3 | 93.2–94.7 | 89.6–91.0 | 91.2–92.4 | 18 ms |
|  | ConvNext (128 × 128) | 93.1–93.5 | 94.6–95.2 | 89.8–90.5 | 91.6–92.2 | 40 ms |
|  | ConvNext (224 × 224) | 93.3–93.4 | 94.7–95.4 | 89.1–90.7 | 91.3–92.4 | 86 ms |

**Table 4.**
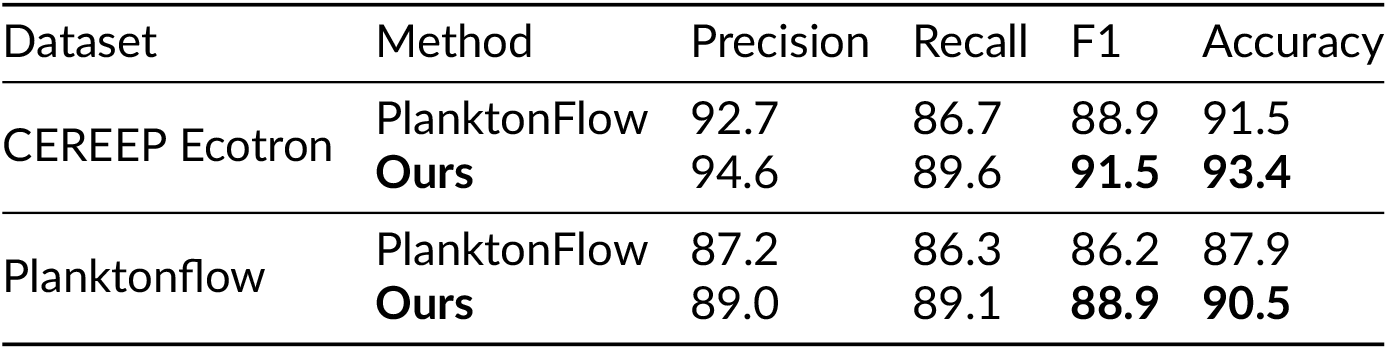
Cross-comparison with the PlanktonFlow method and dataset.

Our results show that the label-cleaning phase improves precision, with V2.1 consistently outperforming V2.0, while recall slightly decreases across all resolutions, resulting in nearly unchanged F1-scores. Accuracy does not increase significantly with V2.1. Moreover, V2.1 exhibits more stable performance across runs, with a smaller gap between the minimum and maximum scores than V2.0. The third observation, which is more surprising, is that image size has no noticeable impact on accuracy. A model trained on smaller images of size 64*×*64 achieves performance comparable to a model trained on images of size 224 *×* 224, while requiring substantially lower training and inference times.

### 4.3. Cross dataset and benchmark

We compare our method with the recently published PlanktonFlow approach (Walter et al., 2025), a deep-learning pipeline for automated plankton image classification that incorporates image preprocessing and offline data augmentation, with a particular focus on rare classes. We conduct experiments to assess the performance of the two approaches. First, the PlanktonFlow method is evaluated on our dataset using the best model and hyperparameters reported by the authors, and making the preprocessing described in the paper. Conversely, our method is tested on the PlanktonFlow dataset. To do so, the raw images from PlanktonFlow are used while preserving the original training and test splits used in the paper. The only preprocessing step retained from PlanktonFlow was the removal of the scale bar; otherwise, our own transformation pipeline was applied. The results of these comparisons are presented section 4.3 with our method slightly outperforming PlanktonFlow both on our dataset and on the PlanktonFlow dataset with higher F1-score and accuracy.

## 5. Conclusion

In this data paper, PlanktonLake-CEREEP, a novel freshwater plankton dataset, is introduced. It includes both large phytoplankton, most zooplankton organisms and a dominant proportion of detritus. Images were acquired with the FlowCam imaging tool and the dataset was designed to be readily integrated into machine learning pipelines. PlanktonLake-CEREEP complements a small number of published and freely available annotated image collections from freshwater plankton Gorzerino et al., 2025; Kyathanahally et al., 2021a; Otake et al., 2024, but differs from those because particular attention was given to label quality through the application of a semi-automatic label correction strategy. For this purpose, a simple classification model based on the ConvNeXt-tiny architecture Liu et al., 2022 is used, designed to be accessible and computationally efficient. Experimental results show that the proposed approach is effective for rapidly identifying potentially suspicious images and that label-cleaning is more efficient by combining complementary methods to find suspicious images. Moreover, the model remains robust to small labeling imperfections, which is likely due to the relatively low number of remaining annotation errors after the first correction process. Even when no significant improvement in overall performance is observed, the proposed approach leads to more consistent results across runs, with lower variance between repeated experiments

Building on this work, we identify several promising research directions to advance freshwater plankton imaging and automatic classification. First, combining already published datasets and image collections from multiple sources such as Ecotaxa could lead to more robust and generalizable classification models of freshwater plankton. Such a combination may require to handle hierarchical and heterogeneous taxonomic annotations, as suggested by Elhamod et al., 2021. Another important direction for the future is out-of-distribution detection, meant to handle, at inference, new or rare species absent from the training data Chen et al., 2025. Future work could also investigate methods for incorporating newly discovered classes after the initial training phase, an area commonly referred to as class-incremental or continual learning Zhou et al., 2024.

## Acknowledgements

We would like to thank Noah-Luc SIMON and Matthieu HINGOUET for their contribution to this work, and Xiaohu Liu for his contribution to the first version of the dataset.

This work was granted access to the HPC resources of IDRIS under the allocation 2025-AD011016996 made by GENCI

## Data, script, code, and supplementary information availability

Data are available online (https://doi.org/10.5281/zenodo.22012196; Déchaumet et al., 2026a)

Script and codes are available online (https://doi.org/10.5281/zenodo.23036077; Déchaumet, 2026)

Supplementary information is available online (https://doi.org/10.5281/zenodo.22940424; Déchaumet et al., 2026b)

